## Supplementary Material for "DNA metabarcoding reveals host-specific communities of arthropods residing in fungal fruit bodies"

### Table of contents

1. Fungal host metadata p. 2
2. Primer pair pilot study p. 3
3. Bioinformatics analyses p. 4
4. Controls and replicates p. 6
5. Rarefaction curves p. 7
6. gNMDS diagnostics p. 8
7. Alpha diversity indices p. 10
8. Univariate testing of H’ p. 11
9. Multivariate testing p. 12
10. Indicator species analysis p. 13

References p. 19

### 1. Fungal host metadata

**Table S1 .** Overview of the eleven wood-decay fungal species and six species-specific fruit body characteristics compiled from the literature. FB size = fruit body size is a factor variable with three levels where 1 = small, 2 = intermediate and 3 = large. Hyphal system complexity relates to toughness and persistence; monomitic fruit bodies are soft and short-lived, while dimitic and trimitic are progressively tougher and long-lived (with the exception of **triabi** which is short-lived). Short-lived is synonymous to annual and long-lived is perennial. Morphology: pileate fruit bodies are voluminous and shelf-like, while resupinate are flat with large surface areas. Hymenophore area ($cm^2$) is the mean spore-producing surface-area. FB mean thickness (cm) is the mean of minimum and maximum fruit body thickness. Sign. OTUs are the number of signficant associations with arthropod OTUs following an indicator species analysis. Number of samples are after sequencing, bioinformatics and data processing, in which low-quality samples were removed. **Postia cyanescens* Miettinen in *Postia caesia* (Schrad.) P. Karst. complex.

| Abb. | Species | FB size | Hyphal system | Persistence | Morphology | Hymenophore area ($cm^2$) | FB mean thickness (cm) | Sign. OTUs | Number of samples |
|---|---|---|---|---|---|---|---|---|---|
| **amylap** | *Amylocystis lapponica* (Romell) Bondartsev & Singer | 3 | Monomitic | Short-lived | Pileate | 53,2 | 30 | 11 | 21 |
| **antser** | *Antrodia serialis* (Fr.) Donk | 2 | Dimitic | Long-lived | Resupinate | 370,6 | 2 | 36 | 23 |
| **fompin** | *Fomitopsis pinicola* (Sw.) P. Karst. | 3 | Trimitic | Long-lived | Pileate | 191,9 | 45 | 12 | 11 |
| **fomros** | *Fomitopsis (Rhodofomes) rosea* (Alb. & Schwein) P. Karst. | 3 | Trimitic | Long-lived | Pileate | 63,9 | 20 | 22 | 12 |
| **glosep** | *Gloeophyllum sepiarium* (Wulfen) P. Karst. | 2 | Trimitic | Long-lived | Pileate | 48,6 | 8.5 | 18 | 15 |
| **phecen** | *Phlebia centrifuga* P. Karst. | 2 | Monomitic | Short-lived | Resupinate | 736 | 2 | 49 | 21 |
| **phefer** | *Phellinus ferrugineofuscus* (P. Karst.) Fiasson & Niemelä | 2 | Dimitic | Long-lived | Resupinate | 494 | 3.5 | 28 | 5 |
| **phenig** | *Phellopilus nigrolimitatus* (Romell) Niemelä, T. Wagner & M. Fisch. | 3 | Dimitic | Long-lived | Resupinate | 218,4 | 7.5 | 33 | 16 |
| **phevit** | *Phellinus (Fuscoporia) viticola* (Schwein.) Donk | 2 | Dimitic | Long-lived | Resupinate | 151,4 | 7.5 | 30 | 14 |
| **poscae** | **Postia cyanescens* Miettinen | 1 | Monomitic | Short-lived | Pileate | 17 | 5 | 7 | 22 |
| **triabi** | *Trichaptum abietinum* (Pers. ex J.F. Gmel.) Ryvarden | 2 | Dimitic | Short-lived | Resupinate | 173 | 3 | 19 | 23 |

### 2. Primer pair pilot study

We tested two primer pairs (9 May 2019)– lCOIintF/jgHCO2198 (313 bp) (Geller et al. 2013, Leray et al. 2013) and LCO1490/230R (300 bp) (Folmer et al. 1994, Gibson et al. 2015) – on 32 fruit body samples representing all eleven wood-decay fungal species and sequenced the amplicons by Sanger sequencing (Fig. S2a). In only six of the Sanger sequences, three non-congruent samples from each pair, there was a metazoan taxon. The BF3/BR2 (421 bp) (Elbrecht et al. 2019) and LCO1490/230R (300 bp) (Folmer et al. 1994, Gibson et al. 2015) primer pairs were tested on seven of the same samples (16 October 2019), including the six with metazoan sequence results, and an additional ten samples (Figure S2b). Based on this optimisation step, we selected the primer pair BF3/BR2 to amplify the arthropods from the fungal fruit bodies gDNA extract, as they had higher amplification success and higher likelihood of fungal identification from BLAST results.

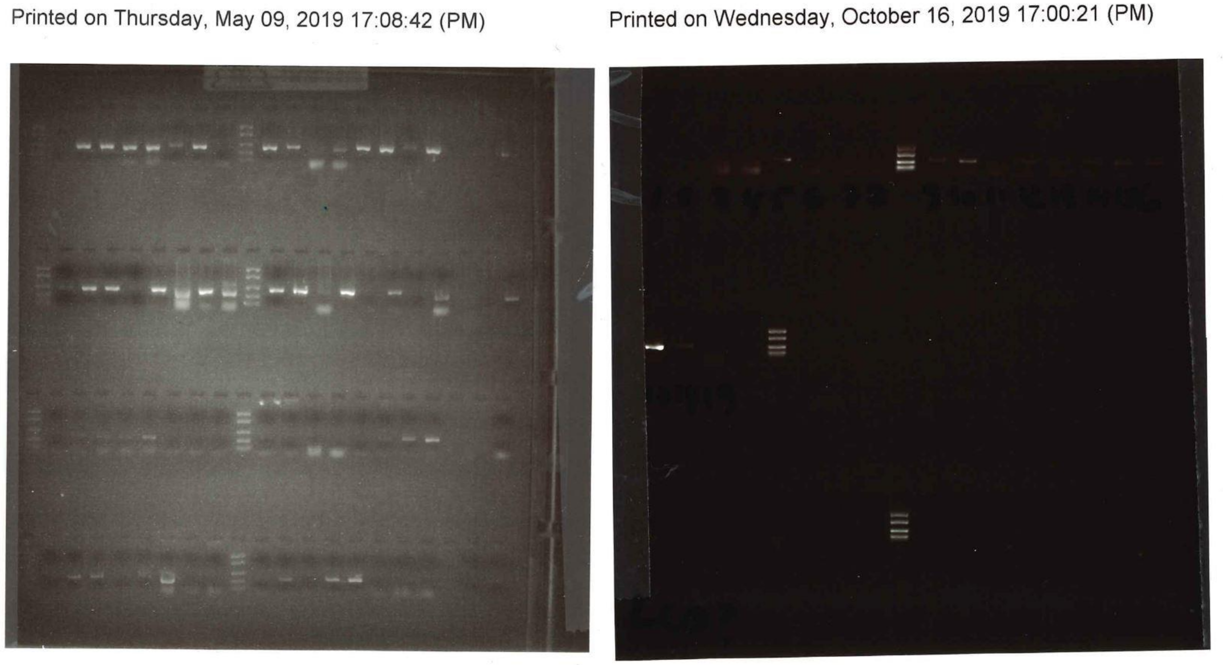

**Figure S2.** Gel electrophoresis screening from initial tests of primer pairs on fungal fruit body samples. (left, **a**) shows mlCOIintF/jgHCO2198 (top lanes) and LCO1490/230R (bottom lanes) from 9. May 2019. (right, **b**) shows BF3/BR2 (top) and LCO1490/230R (bottom) from 16. October 2019.

### 3. Bioinformatics analyses

#### 3.1 Effect of the linker tag

To account for the effect of the linker tag we inserted between the barcode and Illumina adapter, we did two parallel runs of demultiplexing, with and without the linker tag. First, all the barcodes and primers were included for pattern recognition during demultiplexing. Second, the linker tag between the barcode and Illumina adapter was removed. The idea was to investigate the potential effect the linker tag would have on the bioinformatic process. The reverse and forward sequences were demultiplexed with CUTADAPT v 2.7 (Martin 2011) (at least 26 bp overlap, no indels and minimum 100 bp length). The demultiplexing was very similar between the two runs, but slightly more sequences were kept when the linker was removed (Figure S3).

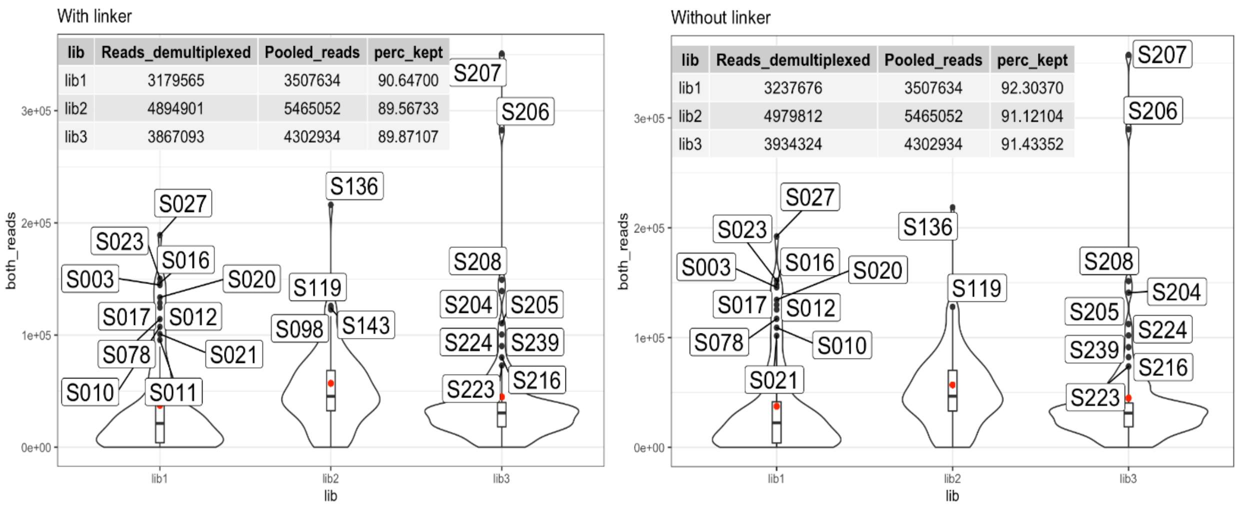

**Figure S3** Sequence depth and proportion of retained sequences after demultiplexing between two parallel bioinformatic runs – with and without linker.

The demultiplexed sequences were then denoised with DADA2 v 1.14 (Callahan et al. 2016) in R v 4.0.3 (Team 2013): low-quality sequences were initially filtered (filterAndTrim: max EE = 2, TruncLen = 260:220), then an error model was estimated from the sequence data (learnErrors: verbose = TRUE, multithread = TRUE) which was used to run the divisive amplicon denoising algorithm (dada: pool = “pseudo”, multithread = TRUE). Then, reverse and forward sequences were merged with overhangs trimmed off and a minimum overlap of 12 bp. Last, chimeras were detected and removed (removeBimeraDenovo: method = “consensus”, multithread = TRUE) to make an amplicon sequence variants (ASV) table. Sequence and ASV distribution across samples, as well as the ASV length, was similar for the two runs, but without the linker tag one more sample was retained. Overall, the effect of the linker tag was minimal for the outcome of the bioinformatic process. We chose to continue processing using the run where the linker tag was removed.

#### 3.1 Effect of clustering

Bioinformatic processing was yet again divided in two parallel runs, one with and one without further clustering of the ASVs into operational taxonomic units (OTUs), to see how clustering would affect taxon richness. Clustering was performed at 97% similarity with VSEARCH (Rognes et al. 2016). Taxonomy was assigned with BLAST+ v 2.8.1 against the BOLD and NCBI databases. Finally, we used the LULU algorithm

(Frøslev et al. 2017) with default settings to correct for potential oversplitting of OTUs. The run without clustering resulted in 13 865 ASVs, against 9316 OTUs when clustering, and we chose to continue with the more conservative, i.e. clustered, dataset to avoid overestimation of biodiversity (Estensmo et al. 2021).

### 4. Controls and replicates

There were three PCR negatives, one per 96-well plate. Two of them were removed during the bioinformatic steps, while the third one contained only 127 sequences which was interpreted as negligible between-sample contamination.

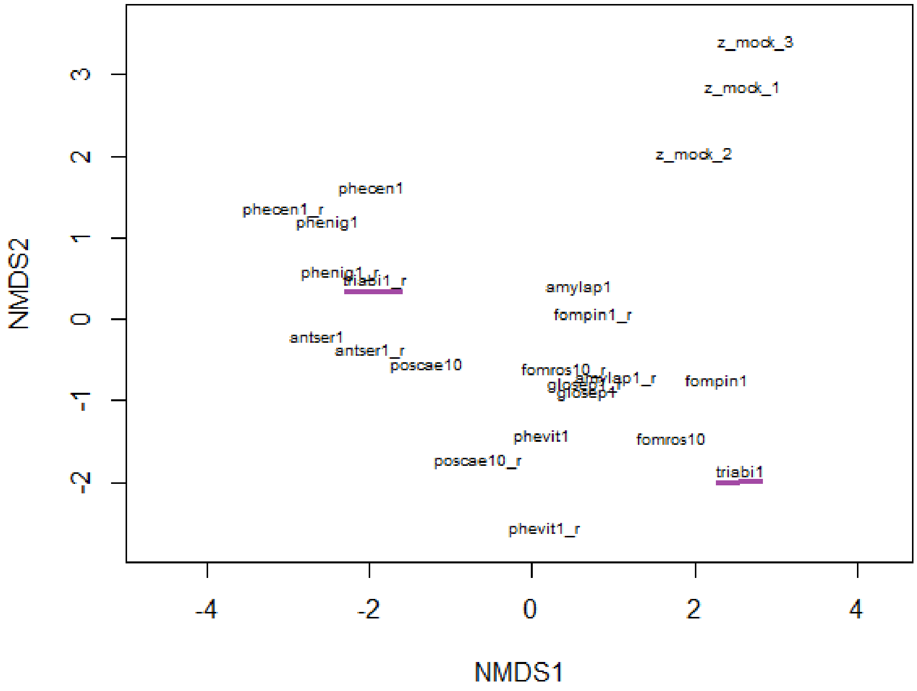

**Figure S4.** Non-metric multidimensional scaling (k=4, stress=0.088) of rarefied PCR and mock replicates. All pairs of technical replicates clustered well except for the pair representing *Trichaptum abietinum* (triabi, underlines in purple).

The mock communities clustered well on the first axis and were dominated by the same OTUs, Lepismatidae sp. – the mock species – and a few species of flies (Figure S4). There were 13 PCR replicates, but three were removed during bioinformatic steps. This resulted in ten pairs, representing all species except for *Phellinus ferrugineofuscum*. Number of sequences between replicate pairs were very similar, except for antser1 (84856 vs 19479), phenig1 (6320 vs 47171) and triabi1 (7951 vs 30). Most pairs clustered well in a gNMDS (Figure S4). The pair that did not cluster had low read numbers in the replicate ("triabi1"; reads 7951/30, OTUs 284/4). For each pair, we removed the sample with the lowest number of reads for further statistical analyses

### 5. Rarefaction curves

To decide subsampling depth of sample rarefaction, we explored the rarefaction curves per host. The 25th percentile (i.e. 1416 sequences per host) was satisfactory as the threshold was clearly below the asymptote saturation for each fungal host (Figure S5).

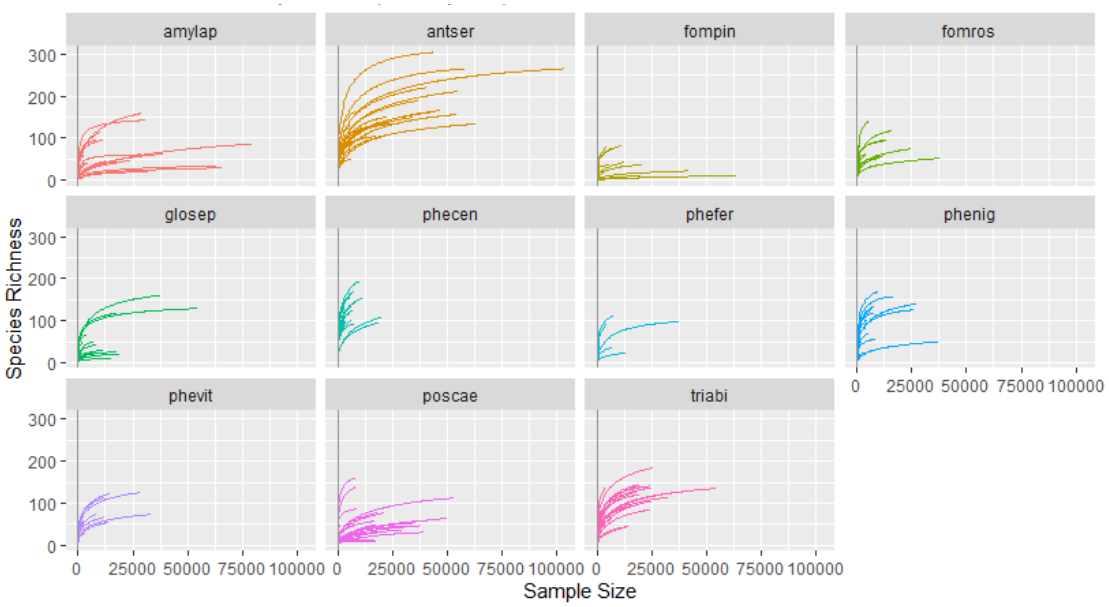

**Figure S5.** Rarefaction curves per fungal host. Vertical lines are drawn at 1416 (the 25th percentile of the full data set) which was the chosen subsampling depth.

### 6. gNMDS diagnostics

After running global non-metric multidimensional scaling (gNMDS) on the arthropod dataset, with settings as suggested by Liu et al. (2008), we compared the two configurations with the lowest stress values with Procrustes analysis (Figure S6.1) and correlation test (999 permutations; $r^2 = 0.99$, $p < 0.01$) (Kent et al. 1979, Peres-Neto and Jackson 2001). After choosing the configuration with the lowest stress values, we inspected it with goodness-of-fit measure and Shepard plot (Figure S6.2). In addition, all four gNMDS axes were congruent with detrended correspondence analysis (DCA) (Hill 1979, Hill and Gauch 1980) ordination axes when compared pair-wise with Kendall’s nonparametric correlations τ (Table S6) (Kendall 1938). The chosen gNMDS configuration is shown in Figure S6.3.

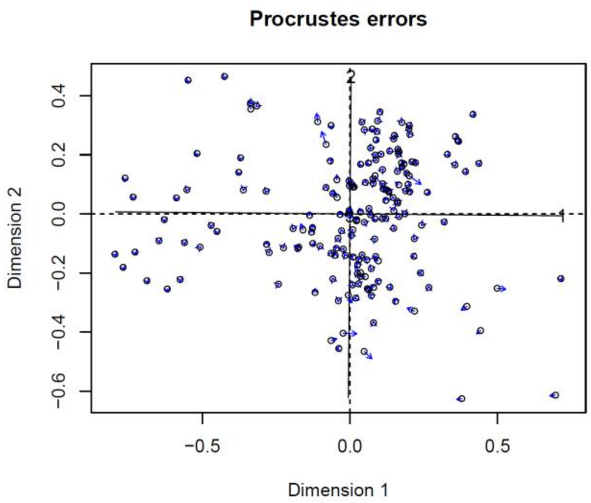

**Figure S6.1.** Procrustes analysis evaluating gNMDS configurations with the lowest and second lowest stress values.

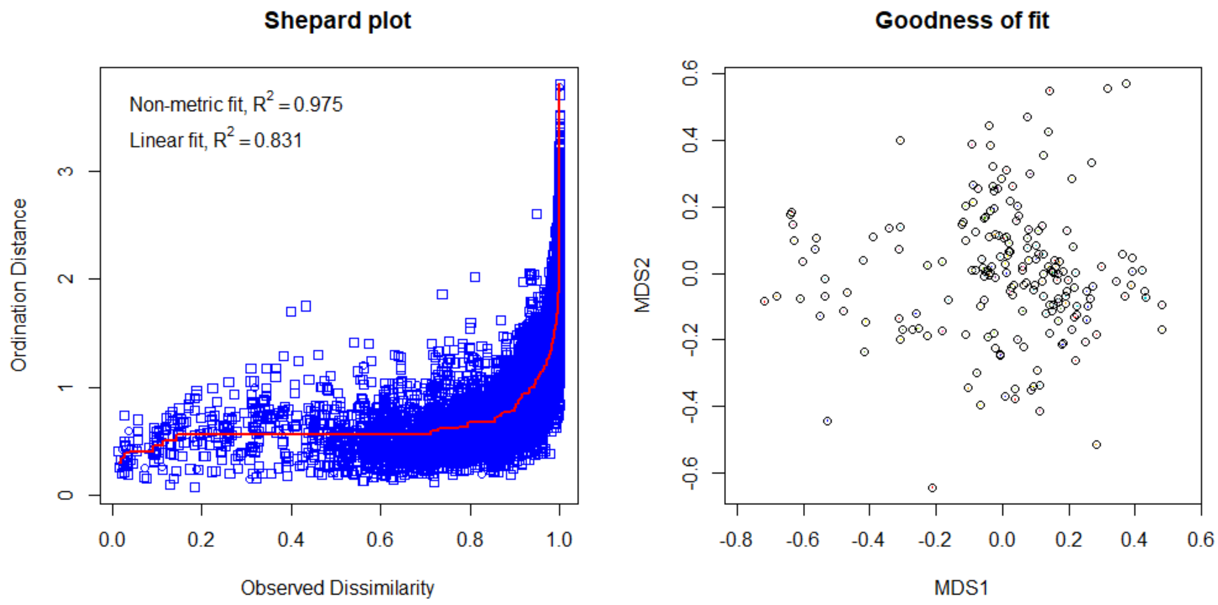

**Figure S6.2.** Shepard plot (left) and goodnes-of-fit measure (right) to evaluate the gNMDS configuration with the lowest stress values.

**Table S6.** Kendall's rank correlation test to compare axes between a detrended correspondence analysis (DCA) and four-dimensional global non-metric multidimensional scaling (gNMDS).

| **DCA axis** | **gNMDS axis** | **tau** | **z** | ***p*** |
|---|---|---|---|---|
| DCA1 | GNMDS1 | -0.49 | -9.7909 | <0.001 |
| DCA1 | GNMDS2 | -0.12 | -2.4793 | 0.013 |
| DCA3 | GNMDS2 | -0.31 | -6.2477 | <0.001 |
| DCA2 | GNMDS3 | 0.19 | 3.7189 | <0.001 |
| DCA4 | GNMDS4 | -0.25 | -4.9808 | <0.001 |

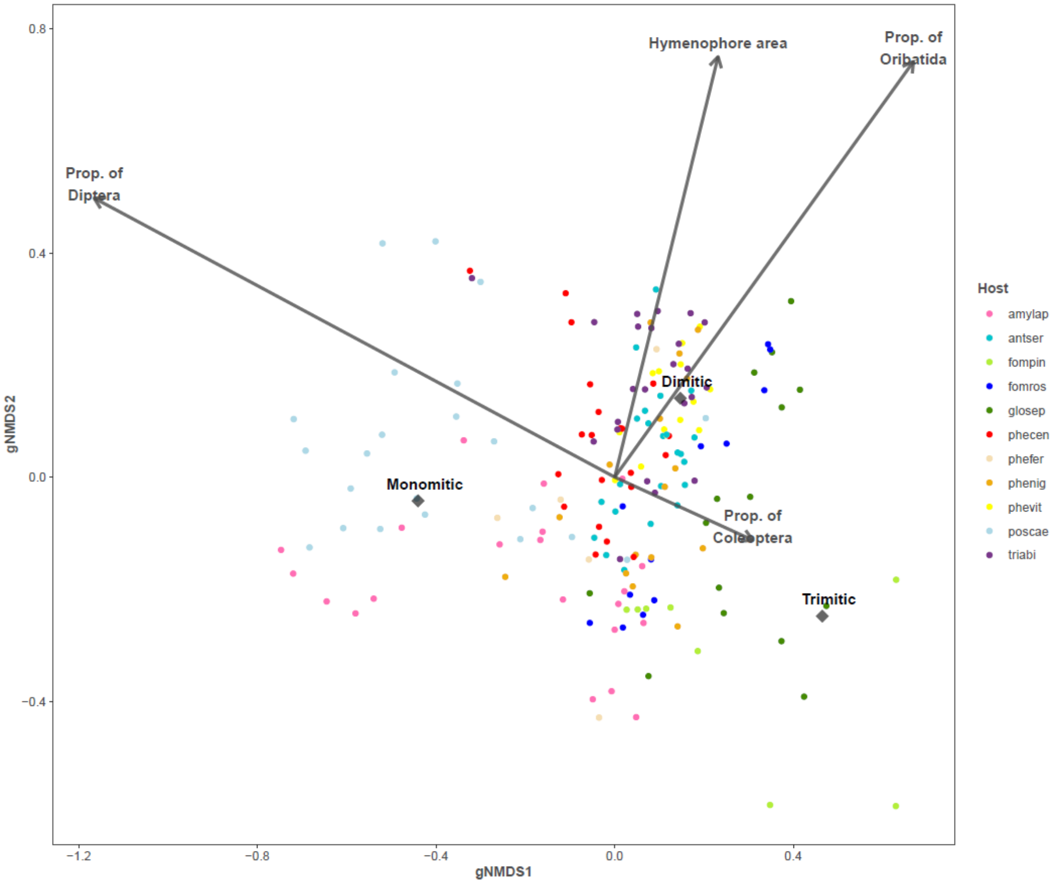

**Figure S6.3. Ordination biplot of arthropod OTUs amplified from representing eleven species of fungal hosts.** The ordination is based on a global non-metric multidimensional scaling (gNMDS; stress = 1.135, k=4) of Bray-Curtis dissimilarities from 180 fungal samples. The axes are sorted by most variation explained and scaled in (half-change) units of compositional turnover. Vectors and centroids are fitted with the envfit function (in package VEGAN) and were significant in multivariate tests. Proportions of arthropod orders were calculated per sample from the data set, not from species scores.

### 7. Alpha diversity indices

The alpha diversity indices Chao1, Shannon and Inverse Simpson were fairly similar when comparing them at the host level (Figure S7). As Inverse Simpson levels were much more compressed and Chao1 were less distinguished between species, we chose Shannon diversity for further analyses.

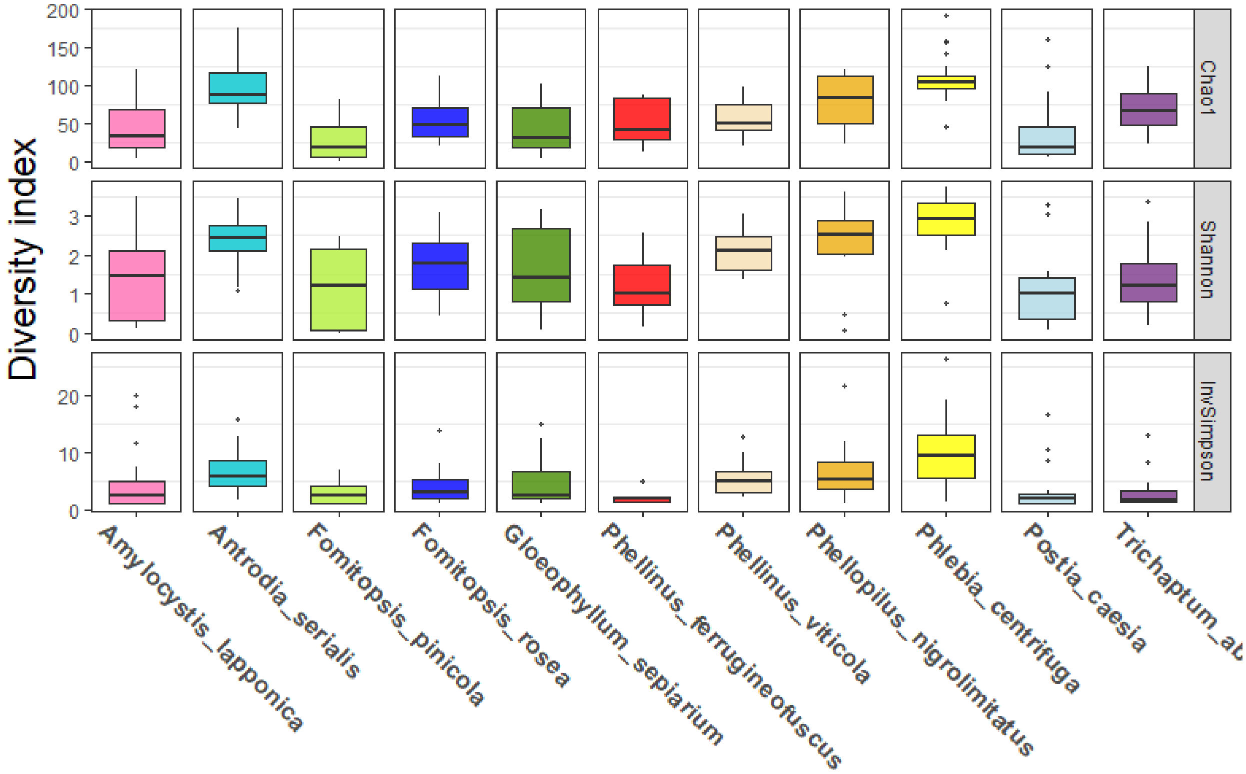

**Figure S7.** Three alpha diversity indices among fungal host species in rarefied arthropod dataset (Chao1, Shannon and Inverse Simpson)

**8. Univariate testing of H'**

**Table S8.** Linear mixed model outputs from five single covariate models explaining the variation in arthropod Shannon diversity with fungal host (eleven species) as random effect. First lines show model intercepts with reference levels of factors in brackets. SE = standard error. Maximum likelihood is optimisation criterion. df (degrees of freedom), t and p calculated from Satterwaite's approximation.

| **Covariate model** | **Estimate** | **SE** | **df** | **t value** | ***p* value** |
|---|---|---|---|---|---|
| **Size (FB size class 1)** | 1,15 | 0,57 | 7,1 | 2 | 0,084 |
| **---------------- class 2** | 0,81 | 0,62 | 7,3 | 1,3 | 0,23 |
| **---------------- class 3** | 0,518 | 0,64 | 7,3 | 0,8 | 0,447 |
| **Persistence (long-lived)** | 1,81 | 0,24 | 9,7 | 7,6 | <0,001 |
| | -0,09 | 0,39 | 8,8 | -0,2 | 0,811 |
| **Hyphal system (monomitic)** | 1,83 | 0,35 | 7,5 | 5,2 | 0,001 |
| **------------------ Dimitic** | 0,09 | 0,45 | 7,8 | 0,2 | 0,852 |
| **------------------ Trimitic** | -0,34 | 0,51 | 8 | -0,7 | 0,526 |
| **Hymenophore surface** | 0,98 | 0,71 | 8,4 | 1,4 | 0,202 |
| | 0,15 | 0,13 | 8,4 | 1,2 | 0,281 |
| **Mean thickness** | 2,01 | 0,23 | 9,2 | 8,9 | <0,001 |
| | -0,02 | 0,01 | 9,6 | -1,5 | 0,161 |

### 9. Multivariate testing

**Table S9.1.** PERMANOVA model output selected from a forward model selection based on the residual sum of squares and subsequent AICc values calculated from an R script by kdyson deposited on GitHub*.

| **Predictors** | Degrees of freedom | Sum of squares | $R^2$ | Mean squares | *F*-ratio | p value |
|---|---|---|---|---|---|---|
| **Hyphal system** | 2 | 5,622 | 0,07208 | 2,8110 | 8,0172 | 0,001 |
| **Fruit body size** | 2 | 4,215 | 0,05404 | 1,2,1074 | 6,0106 | 0,001 |
| **Fruit body thickness (mean)** | 1 | 0,938 | 0,01213 | 0,9385 | 2,6533 | 0,002 |
| **Morphology** | 1 | 0,818 | 0,01049 | 0,0149 | 2,3331 | 0,003 |
| **Hymenophore area** | 1 | 5,184 | 0,06647 | 5,18451 | 14,7864 | 0,001 |
| **Residuals** | 171 | 60,838 | 0,77977 | 0,3537 | | |
| **Total** | 179 | 78 | 1 | | | |

* https://github.com/kdyson/R_Scripts/blob/master/AICc_PERMANOVA.R

**Table S9.2.** Partitioning of variation among sets of environmental variables chosen from a forward model selection with partially (CCA) constrained ordinations. Inertia unit is the summed eigenvalues (if more than one degree of freedom) for the corresponding variable's constrained axes. Inertia is scaled chi-squared. Residual variation is the combined unconstrained axes

| **Environmental variables** | **Inertia unit** | **Fraction of variation explained** | **Degrees of freedom** |
|---|---:|---:|---:|
| Fruit body size | 1,13 | 0,027 | 2 |
| Hyphal system | 1,10 | 0,026 | 2 |
| Hymenophore area | 0,66 | 0,016 | 1 |
| Morphology | 0,58 | 0,014 | 1 |
| Thickness (mean) | 0,52 | 0,012 | 1 |
| Residual | 38,14 | 0,912 | 172 |
| Total | 41,83 | 1,000 | 179 |

### 10. Indicator species analysis

**Table S10.** Significant co-occurrences between arthropod OTUs and eleven fungal hosts from a multi-level pattern analysis based on indicator values. Statistical significance was assessed from permutation analysis and Benjamini-Hochberg adjusted p values with α = 0.05. Metadata and full species names of fungal hosts can be found in Table S1. Taxonomy was assigned with BLAST against the BOLD and NCBI databases.

| otu ID | amylap | antser | fompin | fomros | glosep | phecen | phefer | phenig | phevit | poscae | triabi | Subphylum | Order | Family | |
|---|---|---|---|---|---|---|---|---|---|---|---|---|---|---|---|
| 658 | 0 | 0 | 0 | 0 | 0 | 1 | 0 | 0 | 0 | 0 | 0 | Crustacea | Amphipoda | Amphilochidea | Echinihimedia |
| 4 | 0 | 0 | 0 | 0 | 0 | 0 | 0 | 1 | 0 | 0 | 0 | Arachnida | Araneae | Dionycha | Anyphops |
| 8 | 0 | 1 | 0 | 0 | 0 | 1 | 0 | 0 | 1 | 0 | 0 | Arachnida | Araneae | Linyphiidae | Pacifiphantes |
| 32 | 0 | 0 | 0 | 0 | 0 | 0 | 0 | 1 | 0 | 0 | 0 | Arachnida | Araneae | Nemesiidae | Aname |
| 6 | 0 | 0 | 0 | 0 | 0 | 1 | 1 | 0 | 0 | 0 | 0 | Arachnida | Araneae | Salticidae | Zygoballus |
| 2 | 0 | 0 | 0 | 0 | 1 | 1 | 0 | 0 | 0 | 0 | 1 | Arachnida | Araneae | Theridiidae | Anelosimus |
| 1318 | 0 | 1 | 0 | 0 | 0 | 1 | 0 | 1 | 1 | 0 | 0 | Hexapoda | Archaeognatha | Machilidae | Petridiobius |
| 752 | 0 | 0 | 0 | 0 | 0 | 1 | 0 | 0 | 0 | 0 | 0 | Hexapoda | Coleoptera | Chrysomelidae | Chrysolina |
| 744 | 0 | 1 | 0 | 0 | 0 | 0 | 0 | 1 | 1 | 0 | 0 | Hexapoda | Coleoptera | Dytiscidae | Hydaticus |
| 755 | 0 | 0 | 0 | 0 | 0 | 0 | 0 | 1 | 0 | 0 | 0 | Hexapoda | Coleoptera | Elateridae | Limoniscus |
| 735 | 0 | 0 | 0 | 0 | 1 | 0 | 0 | 0 | 0 | 0 | 0 | Hexapoda | Coleoptera | Mordellidae | Curtimorda |
| 737 | 0 | 1 | 0 | 1 | 1 | 1 | 0 | 0 | 1 | 0 | 1 | Hexapoda | Coleoptera | Scarabaeidae | Kheper |
| 741 | 0 | 0 | 0 | 0 | 0 | 1 | 0 | 0 | 0 | 0 | 0 | Hexapoda | Coleoptera | Scarabaeidae | Scarabaeus |
| 819 | 0 | 1 | 0 | 0 | 0 | 0 | 0 | 0 | 1 | 0 | 0 | Hexapoda | Collembola | | |
| 686 | 0 | 0 | 0 | 0 | 0 | 0 | 0 | 1 | 0 | 0 | 0 | Crustacea | Copepoda | | |
| 692 | 0 | 0 | 0 | 0 | 0 | 0 | 0 | 1 | 0 | 0 | 0 | Crustacea | Decapoda | Pleocyemata | Calcinus |
| 1358 | 0 | 0 | 0 | 0 | 0 | 0 | 0 | 1 | 0 | 0 | 0 | Myriapoda | Diplopoda | Lithobiidae | Lithobius |

| | | | | | | | | | | | | | | | |
|---|---|---|---|---|---|---|---|---|---|---|---|---|---|---|---|
| 879 | 0 | 1 | 0 | 1 | 0 | 1 | 0 | 0 | 1 | 0 | 1 | Hexapoda | Diptera | Agromyzidae | |
| 851 | 1 | 0 | 0 | 0 | 0 | 0 | 1 | 0 | 0 | 1 | 0 | Hexapoda | Diptera | Bolitophilidae | |
| 854 | 0 | 0 | 0 | 0 | 0 | 0 | 0 | 0 | 0 | 1 | 0 | Hexapoda | Diptera | Bolitophilidae | |
| 870 | 1 | 0 | 0 | 0 | 0 | 0 | 0 | 0 | 0 | 1 | 0 | Hexapoda | Diptera | Bolitophilidae | |
| 871 | 1 | 0 | 0 | 0 | 0 | 0 | 0 | 0 | 0 | 1 | 0 | Hexapoda | Diptera | Bolitophilidae | |
| 853 | 1 | 0 | 0 | 0 | 0 | 0 | 1 | 0 | 0 | 0 | 0 | Hexapoda | Diptera | Cecidomyiidae | |
| 860 | 0 | 0 | 0 | 0 | 1 | 0 | 0 | 0 | 0 | 0 | 0 | Hexapoda | Diptera | Cecidomyiidae | |
| 863 | 0 | 0 | 1 | 0 | 0 | 0 | 0 | 0 | 0 | 0 | 0 | Hexapoda | Diptera | Cecidomyiidae | Lestodiplosis |
| 865 | 0 | 0 | 0 | 0 | 1 | 0 | 0 | 0 | 0 | 0 | 0 | Hexapoda | Diptera | Cecidomyiidae | |
| 852 | 0 | 0 | 0 | 0 | 0 | 0 | 0 | 0 | 0 | 0 | 1 | Hexapoda | Diptera | Chironomidae | |
| 883 | 0 | 0 | 0 | 0 | 0 | 1 | 0 | 0 | 0 | 0 | 0 | Hexapoda | Diptera | Limoniidae | Metalimnobia |
| 928 | 0 | 0 | 0 | 0 | 0 | 1 | 0 | 0 | 0 | 0 | 1 | Hexapoda | Diptera | Muscidae | Limnophora |
| 855 | 0 | 0 | 0 | 0 | 0 | 0 | 0 | 0 | 0 | 1 | 0 | Hexapoda | Diptera | Mycetophilidae | Mycetophila |
| 891 | 0 | 0 | 0 | 0 | 0 | 0 | 0 | 0 | 0 | 1 | 0 | Hexapoda | Diptera | Mycetophilidae | Mycetophila |
| 919 | 0 | 0 | 0 | 0 | 0 | 0 | 0 | 1 | 0 | 0 | 0 | Hexapoda | Diptera | Mycetophilidae | Sciophila |
| 942 | 0 | 0 | 0 | 1 | 0 | 0 | 0 | 0 | 0 | 0 | 0 | Hexapoda | Diptera | Mycetophilidae | Dynatosoma |
| 961 | 0 | 0 | 0 | 0 | 0 | 0 | 1 | 0 | 0 | 0 | 0 | Hexapoda | Diptera | Mydidae | Rhaphiomidas |
| 916 | 0 | 0 | 0 | 0 | 0 | 0 | 0 | 1 | 0 | 0 | 0 | Hexapoda | Diptera | Pipunculidae | Tomosvaryella |
| 856 | 0 | 1 | 0 | 0 | 0 | 0 | 0 | 0 | 0 | 0 | 0 | Hexapoda | Diptera | Sciaridae | |
| 864 | 0 | 0 | 0 | 0 | 0 | 1 | 0 | 1 | 0 | 0 | 0 | Hexapoda | Diptera | Sciaridae | |
| 892 | 0 | 0 | 0 | 1 | 0 | 0 | 0 | 0 | 0 | 0 | 0 | Hexapoda | Diptera | Sciaridae | |

| | | | | | | | | | | | | | | | |
|---|---|---|---|---|---|---|---|---|---|---|---|---|---|---|---|
| 868 | 0 | 0 | 0 | 0 | 0 | 1 | 1 | 0 | 0 | 0 | 0 | Hexapoda | Diptera | Tachinidae | Mactomyia |
| 1124 | 0 | 0 | 0 | 0 | 0 | 1 | 0 | 0 | 1 | 0 | 0 | Hexapoda | Hemiptera | Aphididae | |
| 1130 | 0 | 0 | 0 | 0 | 0 | 0 | 0 | 1 | 0 | 0 | 0 | Hexapoda | Hemiptera | Notonectidae | Notonecta |
| 1120 | 0 | 1 | 0 | 0 | 0 | 1 | 1 | 0 | 1 | 0 | 1 | Hexapoda | Hemiptera | | |
| 1142 | 0 | 0 | 0 | 0 | 1 | 0 | 0 | 0 | 0 | 0 | 0 | Hexapoda | Hemiptera | | |
| 1198 | 0 | 1 | 0 | 0 | 0 | 1 | 0 | 1 | 0 | 0 | 0 | Hexapoda | Hymenoptera | Braconidae | Promicrogaster |
| 1201 | 0 | 0 | 0 | 0 | 0 | 1 | 0 | 0 | 0 | 0 | 0 | Hexapoda | Hymenoptera | Braconidae | Diolcogaster |
| 1197 | 0 | 1 | 0 | 0 | 0 | 0 | 0 | 0 | 1 | 1 | 0 | Hexapoda | Hymenoptera | Tenthredinidae | Dolerus_vestigialis |
| 1254 | 0 | 1 | 0 | 0 | 0 | 1 | 0 | 0 | 1 | 0 | 0 | Hexapoda | Lepidoptera | Noctuidae | Mocis |
| 1256 | 0 | 0 | 0 | 1 | 0 | 0 | 0 | 0 | 0 | 0 | 0 | Hexapoda | Lepidoptera | Noctuidae | Apamea |
| 1260 | 0 | 0 | 0 | 1 | 0 | 0 | 0 | 0 | 0 | 0 | 0 | Hexapoda | Lepidoptera | Tineidae | Agnathosia_mendicella |
| 103 | 0 | 0 | 1 | 0 | 0 | 0 | 0 | 0 | 0 | 0 | 0 | Arachnida | Mesostigmata | Blattisociidae | Lasioseius |
| 106 | 0 | 0 | 0 | 0 | 0 | 0 | 0 | 1 | 0 | 0 | 0 | Arachnida | Mesostigmata | Phytoseiidae | Euseius |
| 129 | 0 | 0 | 1 | 0 | 1 | 1 | 0 | 0 | 0 | 0 | 1 | Arachnida | Oribatida | Carabodidae | Carabodes |
| 131 | 0 | 1 | 0 | 0 | 0 | 1 | 0 | 0 | 0 | 0 | 0 | Arachnida | Oribatida | Ceratoppiidae | Ceratoppia |
| 125 | 0 | 1 | 0 | 1 | 1 | 1 | 1 | 1 | 1 | 0 | 1 | Arachnida | Oribatida | Oppiidae | Oppia |
| 139 | 0 | 0 | 0 | 0 | 0 | 0 | 1 | 0 | 0 | 0 | 0 | Arachnida | Oribatida | Phthiracaridae | Phthiracarus |
| 126 | 0 | 1 | 0 | 0 | 0 | 1 | 0 | 1 | 1 | 0 | 1 | Arachnida | Oribatida | Suctobelbidae | |
| 220 | 0 | 0 | 0 | 0 | 0 | 0 | 0 | 0 | 1 | 0 | 0 | Arachnida | Prostigmata | Ereynetidae | |
| 212 | 0 | 0 | 0 | 1 | 0 | 0 | 0 | 0 | 0 | 0 | 0 | Arachnida | Prostigmata | Tarsonemidae | |

| | | | | | | | | | | | | |
|---|---|---|---|---|---|---|---|---|---|---|---|---|
| 275 | 0 | 1 | 0 | 1 | 0 | 1 | 0 | 0 | 1 | 0 | 1 | Arachnida |
| 276 | 1 | 1 | 0 | 1 | 0 | 1 | 1 | 0 | 1 | 0 | 0 | Arachnida |
| 277 | 1 | 1 | 0 | 0 | 0 | 1 | 1 | 0 | 1 | 0 | 0 | Arachnida |
| 278 | 0 | 1 | 0 | 1 | 0 | 1 | 0 | 0 | 1 | 0 | 0 | Arachnida |
| 280 | 0 | 1 | 0 | 1 | 1 | 1 | 1 | 1 | 1 | 0 | 0 | Arachnida |
| 281 | 0 | 1 | 0 | 0 | 0 | 1 | 1 | 0 | 1 | 0 | 1 | Arachnida |
| 282 | 0 | 1 | 0 | 1 | 1 | 1 | 1 | 0 | 0 | 0 | 0 | Arachnida |
| 283 | 0 | 1 | 0 | 0 | 0 | 1 | 1 | 1 | 1 | 0 | 0 | Arachnida |
| 284 | 0 | 0 | 0 | 0 | 0 | 1 | 0 | 1 | 0 | 0 | 0 | Arachnida |
| 289 | 0 | 0 | 0 | 0 | 0 | 1 | 1 | 0 | 0 | 0 | 0 | Arachnida |
| 290 | 0 | 1 | 0 | 0 | 0 | 1 | 1 | 0 | 1 | 0 | 1 | Arachnida |
| 293 | 0 | 0 | 0 | 0 | 0 | 0 | 0 | 0 | 0 | 0 | 1 | Arachnida |
| 294 | 0 | 1 | 0 | 0 | 0 | 0 | 0 | 1 | 0 | 0 | 0 | Arachnida |
| 295 | 0 | 0 | 0 | 0 | 0 | 0 | 1 | 0 | 1 | 0 | 0 | Arachnida |
| 303 | 0 | 0 | 1 | 0 | 1 | 1 | 0 | 0 | 0 | 0 | 0 | Arachnida |
| 312 | 0 | 0 | 0 | 0 | 0 | 1 | 0 | 0 | 0 | 0 | 0 | Arachnida |
| 314 | 0 | 0 | 0 | 0 | 0 | 1 | 0 | 0 | 0 | 0 | 1 | Arachnida |
| 315 | 0 | 0 | 0 | 0 | 0 | 1 | 0 | 0 | 0 | 0 | 0 | Arachnida |
| 316 | 0 | 0 | 0 | 0 | 0 | 1 | 1 | 0 | 0 | 0 | 0 | Arachnida |
| 318 | 0 | 0 | 0 | 0 | 0 | 1 | 0 | 0 | 0 | 0 | 0 | Arachnida |
| 327 | 0 | 0 | 0 | 0 | 0 | 0 | 0 | 0 | 0 | 0 | 1 | Arachnida |

| | | | | | | | | | | | | |
|---|---|---|---|---|---|---|---|---|---|---|---|---|
| 329 | 0 | 0 | 0 | 0 | 0 | 1 | 0 | 0 | 0 | 0 | 0 | Arachnida |
| 330 | 0 | 0 | 0 | 0 | 0 | 0 | 0 | 1 | 0 | 0 | 0 | Arachnida |
| 332 | 0 | 0 | 0 | 0 | 0 | 0 | 0 | 1 | 0 | 0 | 0 | Arachnida |
| 333 | 0 | 0 | 0 | 0 | 0 | 0 | 0 | 0 | 0 | 0 | 1 | Arachnida |
| 342 | 0 | 0 | 0 | 0 | 0 | 0 | 0 | 1 | 0 | 0 | 0 | Arachnida |
| 356 | 0 | 0 | 0 | 0 | 0 | 0 | 1 | 0 | 0 | 0 | 1 | Arachnida |
| 360 | 0 | 0 | 0 | 0 | 0 | 1 | 0 | 0 | 0 | 0 | 0 | Arachnida |
| 370 | 0 | 0 | 0 | 0 | 0 | 0 | 0 | 1 | 0 | 0 | 0 | Arachnida |
| 380 | 0 | 0 | 0 | 0 | 0 | 1 | 0 | 0 | 0 | 0 | 0 | Arachnida |
| 1367 | 0 | 0 | 0 | 0 | 0 | 0 | 1 | 0 | 0 | 0 | 0 | unk. Arthropoda |
| 1368 | 0 | 1 | 0 | 1 | 0 | 0 | 0 | 1 | 1 | 0 | 0 | unk. Arthropoda |
| 1369 | 1 | 0 | 0 | 1 | 0 | 0 | 0 | 0 | 0 | 0 | 0 | unk. Arthropoda |
| 1370 | 0 | 0 | 0 | 0 | 1 | 0 | 0 | 0 | 0 | 0 | 0 | unk. Arthropoda |
| 1371 | 1 | 1 | 1 | 1 | 0 | 1 | 1 | 0 | 1 | 0 | 0 | unk. Arthropoda |
| 1372 | 1 | 0 | 1 | 0 | 0 | 0 | 0 | 1 | 0 | 0 | 0 | unk. Arthropoda |
| 1374 | 0 | 1 | 1 | 0 | 0 | 0 | 0 | 0 | 0 | 0 | 0 | unk. Arthropoda |
| 1376 | 1 | 1 | 1 | 1 | 1 | 1 | 1 | 1 | 1 | 0 | 1 | unk. Arthropoda |
| 1378 | 0 | 1 | 1 | 1 | 1 | 1 | 1 | 0 | 0 | 0 | 0 | unk. Arthropoda |
| 1379 | 0 | 1 | 0 | 0 | 0 | 0 | 1 | 0 | 0 | 0 | 0 | unk. Arthropoda |
| 1381 | 0 | 1 | 0 | 0 | 1 | 0 | 0 | 0 | 0 | 0 | 0 | unk. Arthropoda |
| 1383 | 0 | 0 | 0 | 0 | 0 | 0 | 1 | 1 | 0 | 0 | 0 | unk. Arthropoda |
| 1384 | 0 | 0 | 0 | 1 | 1 | 0 | 0 | 0 | 0 | 0 | 0 | unk. Arthropoda |
| 1389 | 1 | 0 | 1 | 0 | 0 | 0 | 0 | 0 | 0 | 0 | 0 | unk. Arthropoda |
| 1392 | 0 | 0 | 0 | 1 | 0 | 1 | 0 | 0 | 0 | 0 | 0 | unk. Arthropoda |
| 1395 | 0 | 0 | 0 | 1 | 0 | 1 | 1 | 0 | 0 | 0 | 0 | unk. Arthropoda |
| 1398 | 0 | 0 | 0 | 0 | 1 | 1 | 0 | 0 | 0 | 0 | 1 | unk. Arthropoda |
| 1400 | 0 | 0 | 0 | 0 | 0 | 0 | 0 | 1 | 1 | 0 | 0 | unk. Arthropoda |
| 1403 | 0 | 1 | 0 | 0 | 0 | 0 | 1 | 0 | 0 | 0 | 0 | unk. Arthropoda |

| | | | | | | | | | | | | |
|---|---|---|---|---|---|---|---|---|---|---|---|---|
| 1405 | 0 | 1 | 0 | 0 | 0 | 0 | 0 | 1 | 0 | 0 | 0 | unk. Arthropoda |
| 1406 | 0 | 0 | 1 | 0 | 0 | 0 | 0 | 1 | 1 | 0 | 0 | unk. Arthropoda |
| 1412 | 0 | 1 | 0 | 1 | 0 | 0 | 0 | 0 | 0 | 0 | 0 | unk. Arthropoda |
| 1415 | 0 | 1 | 0 | 0 | 0 | 0 | 0 | 1 | 1 | 0 | 0 | unk. Arthropoda |
| 1432 | 0 | 0 | 0 | 0 | 0 | 0 | 0 | 1 | 0 | 0 | 0 | unk. Arthropoda |
| 1449 | 0 | 0 | 1 | 0 | 0 | 0 | 0 | 0 | 0 | 0 | 0 | unk. Arthropoda |
| 1478 | 0 | 0 | 0 | 0 | 1 | 0 | 0 | 0 | 0 | 0 | 0 | unk. Arthropoda |
| 1484 | 0 | 0 | 0 | 0 | 0 | 0 | 0 | 0 | 1 | 0 | 0 | unk. Arthropoda |
| 1503 | 0 | 1 | 0 | 0 | 0 | 1 | 1 | 0 | 0 | 0 | 0 | unk. Arthropoda |
| 1549 | 0 | 0 | 0 | 0 | 0 | 0 | 0 | 0 | 1 | 0 | 0 | unk. Arthropoda |
